# Intra-Articular Delivery of an Indoleamine 2,3-Dioxygenase Galectin-3 Fusion Protein for Osteoarthritis Treatment in Male Lewis Rats

**DOI:** 10.1101/2021.07.21.453247

**Authors:** Brittany D. Partain, Evelyn Bracho-Sanchez, Shaheen A. Farhadi, Elena G. Yarmola, Benjamin G. Keselowsky, Gregory A. Hudalla, Kyle D. Allen

## Abstract

**Objective:** Controlling joint inflammation can improve osteoarthritis (OA) symptoms; however, current treatments often fail to provide long-term effects. We have developed an indoleamine 2,3-dioxygenase and galectin-3 fusion protein (IDO-Gal3). IDO converts tryptophan to kynurenines, directing the local environment toward an anti-inflammatory state; Gal3 binds carbohydrates and extends IDO’s joint residence time. In this study, we evaluated IDO-Gal3’s ability to alter OA-associated inflammation and pain-related behaviors in a rat model of established knee OA.

**Methods:** Joint residence was first evaluated with an analog Gal3 fusion protein (NanoLuc™ and Gal3, NL-Gal3) that produces luminescence from furimazine. OA was induced in male Lewis rats via a medial collateral ligament and medial meniscus transection (MCLT+MMT). At 8 weeks, NL or NL-Gal3 were injected intra-articularly (n=8 per group), and bioluminescence was tracked for 4 weeks. Next, IDO-Gal3’s ability to modulate OA pain and inflammation was assessed. Again, OA was induced via MCLT+MMT in male Lewis rats, with IDO-Gal3 or saline injected into OA-affected knees at 8 weeks post-surgery (n=7 per group). Gait and tactile sensitivity were then assessed weekly. At 12 weeks, intra-articular levels of IL6, CCL2, and CTXII were assessed.

**Results:** The Gal3 fusion increased joint residence in OA and contralateral knees (p<0.0001). In OA-affected animals, IDO-Gal3 improved tactile sensitivity (p=0.002), increased walking velocities (p≤0.033), and improved vertical ground reaction forces (p≤0.04). Finally, IDO-Gal3 decreased intra-articular IL6 levels within the OA-affected joint (p=0.0025).

**Conclusion:** Intra-articular IDO-Gal3 delivery provided long-term modulation of joint inflammation and pain-related behaviors in rats with established OA.

## Introduction

Osteoarthritis (OA) is associated with chronic local, low grade inflammation (1). Controlling joint inflammation is important, especially if control of inflammation leads to improvement of OA symptoms. Current treatments for OA include oral non-steroidal anti-inflammatory drugs (NSAIDs) and intra-articular injections of corticosteroids or hyaluronic acid (2). Although intra-articular injections increase the local concentration of an anti-inflammatory drug and decrease off target effects, the efficacy of this approach is often hindered by rapid joint clearance (3). Short joint residence times typically necessitate repeated injections, which can lead to increased risk of infection and chondrocyte death (4–6).

Multiple intra-articular delivery strategies aim to extend joint residence times (7). For example, therapeutics are commonly encapsulated in carriers that extend joint residence; however, harsh encapsulation conditions can reduce a drug’s bioactivity (8,9). Alternatively, conjugating protein-based drugs to synthetic particles can extend joint residence, improve bioactivity, and limit degradation (10); however, determining an effective dose to circumvent joint clearance and maintain efficacy has proven difficult (11). In this regard, enzyme-based therapeutics may offer an advantage (12). With the catalytic capabilities of enzymes, target molecules can be continually and locally produced, as long as the enzyme’s substrate is available. This decreases the necessary dose of the enzyme, and conjugating enzymes to anchoring molecules can further enhance bioactivity and extend joint residence.

To provide long-lasting local control of inflammation, we developed an indoleamine 2,3-dioxygenase galectin-3 fusion protein (IDO-Gal3). IDO is an immunomodulatory enzyme that converts tryptophan into kynurenines (13–16). By locally degrading tryptophan, IDO induces effector T cell apoptosis, promotes regulatory T cell proliferation, modulates macrophage differentiation toward an anti-inflammatory phenotype, and maintains an immature phenotype on dendritic cells that results in suppression of antigen-specific T cell proliferation (15,17–19). Gal3 binds to β-galactoside glycoconjugates and glycosaminoglycans, and these sites are prevalent in multiple joint tissues. Via this binding, Gal3 can extend an enzyme’s local residence time (20–22). In this work, IDO-Gal3’s ability to alter OA-associated inflammation and pain-related behaviors was evaluated in a rat model of established knee OA.

## Methods

### Experimental Designs

Animal procedures were approved by the University of Florida’s Institutional Animal Care and Use Committee. In our first experiment, joint retention of a Gal3 fusion protein was assessed (Supplemental Fig. 1). Here, 16 male Lewis rats (250 g) were acquired from Charles River Laboratories (Wilmington, MA, USA) and acclimated to the University of Florida for 1 week. Rats then underwent a medial collateral ligament and medial meniscus transection surgery (MCLT+MMT). At 8 weeks post-surgery, background luminescence was measured for each rat (1 sec exposure, field of view D, PerkinElmer IVIS, Waltham, MA, USA). Both knees were then injected with either 50 μL NanoLuc™ Luciferase (NL, n=8, 3.27 μM) or a NanoLuc™ Luciferase galectin-3 fusion protein (NL-Gal3, n=8, 3.27 μM). For luminescence imaging, knees were injected with 50 μL furimazine (1:50 dilution in PBS, Nano-Glo™, PRN1120, Promega, Madison, WI, USA), then assessed with the above IVIS settings. Luminescence imaging was repeated 1, 2, 4, 8, 12, 16, 20, 24, and 28 days after NL or NL-Gal 3 injection. On day 28, rats were euthanized via exsanguination under deep anesthesia, and the patella, tibia, femur, and meniscus were dissected, preserving the synovium and fat pad connected to each bone. Tissue samples were placed in a 24 well plate and incubated with diluted furimazine for 1 min at 25°C. After incubation, tissues were removed, and luminescence was measured using the autoexposure setting in field of view C.

In our second experiment, we evaluated IDO’s ability to modulate OA-related inflammation and symptoms (Supplemental Fig. 2). Here, 16 additional male Lewis rats (250 g, 3 months) were acquired from Charles River Laboratories and acclimated to the University of Florida for 1 week. Rats then underwent baseline behavioral testing followed by MCLT+MMT surgery; one rat was euthanized due to surgical complications. Post-surgery, rodent gait was assessed at 3, 5, and 7 weeks and tactile sensitivity was assessed weekly (except for week 6 due to university closure). At 8 weeks post-surgery, rats received 30 μL injections of either saline (n=7) or IDO-Gal3 (n=8, 33 pg/μL) in the MCLT+MMT knee. Tactile sensitivity testing was conducted the day after injection, then weekly out to 11 weeks post-surgery. Gait testing was performed 2 days after injection, then weekly out to 11 weeks post-surgery. Rats were then euthanized via exsanguination under deep anesthesia. Immediately post-mortem, IL6, CCL2, and CTXII were assessed in both knees using magnetic capture. Here, saline samples from a contralateral and operated limb were lost due to insufficient collection of magnetic beads. Following magnetic capture, knees were processed for histology. In histology, one IDO-Gal3 treated rat showed an intact medial meniscus (failure to fully transect the meniscus); this animal was removed from the analysis. Thus, the final datasets are n=6-7 for saline treatment and n=7 for IDO-Gal3 treatment. Statistical analyses are described for each method below, with analyses performed at *α*=0.05 in R Studio.

### Detailed Methods

#### Protein Production

Genes encoding NL-Gal3 (IDT, Coralville, IA, USA) and recombinant human Gal3 (OriGene, Rockville, MD, USA) were inserted into pET-21d(+) vectors between the Ncol and Xhol sites, then transformed into One Shot™ TOP10 Chemically Competent E. coli (ThermoFisher, Waltham, MA, USA). NL was amplified from the NL-Gal3 gene using PCR and mutated to have Ncol and Xhol flank the genes using the following primers: 5’-CGC CTC GAG CGC CAG AAT GCG TT-3’ and 5’-GCT TAG CCA TGG CGG TCT TCA CAC TCG AAG-3’. PCR products were digested using Ncol and Xhol and re-inserted into pET-21d(+) vectors. Genes encoding IDO were generously provided by Dr. Carlos Gonzalez’s Laboratory at the University of Florida. Amplification and insertion of the BamHI restriction site was performed using the following primers: 5’-CAG CTA CCA TGG CAC ACG CTA TGG AAA-3’ and GAG AAC 5’-GGA TCC ACC TTC CTT CAA AAG-3’. The NL-Gal3 gene was digested with Ncol and BamHI to remove the NL gene and insert the IDO gene. After overnight treatment on ampicillin (100 μg/mL) doped LB/agar plates (37°C), positive clones were used to inoculate 5 mL of LB broth containing ampicillin (100 μg/mL). Cultures were grown overnight on an orbital shaker (37°C, 220 rpm), with plasmids isolated using a Plasmid Miniprep Kit (Qiagen, Hilden, Germany).

Origami™ B (DE3) E. coli (Novagen, Madison, WI, USA) were transformed with pET-21d-Gal3, pET-21d-NL, pET-21d-NL-Gal3, or pET-21d-IDO-Gal3 vectors and selected on ampicillin (100 μg/mL) and kanamycin A (50 μg/mL) doped LB/agar plates overnight (37°C). Positive clones were used to inoculate 5 mL of LB broth containing ampicillin (100 μg/mL) and kanamycin A (50 μg/mL) and grown overnight (37°C, 220 rpm). Cultures were expanded in 1 L 2xTY broth containing ampicillin (100 μg/mL) and kanamycin A (50 μg/mL) and grown on an orbital shaker (37°C, 225 rpm) until the optical density reached 0.6-0.8 (λ=600). Protein expression was induced via 0.5 mM isopropyl β-D-1-thiogalactopyranoside (ThermoFisher, Waltham, MA, USA) for 18 hrs (18°C, 225 rpm). Bacteria were pelleted via centrifugation, washed with PBS, and incubated for 20 mins with B-PER™ bacterial protein extraction reagent (ThermoFisher, Waltham, MA, USA), 1 Pierce protease inhibitor tablet (ThermoFisher, Waltham, MA, USA), 300 units DNAse I from bovine pancreas (MilliporeSigma, Burlington, MA, USA), and 100 μg lysozyme (MilliporeSigma, Burlington, MA, USA). Lysed bacteria were centrifuged, and supernatant was loaded into columns containing HisPur™ Cobalt Superflow Agarose (ThermoFisher, Waltham, MA, USA). Proteins were eluted using an imidazole gradient (0-250 mM), then centrifuged on Amicon Ultra Centrifugal Filters with a 10 kDa cutoff (MilliporeSigma, Burlington, MA, USA). A second purification step was performed for IDO-Gal3 using size exclusion chromatography in an ÄKTA Pure Protein Purification System (GE Life Sciences, Marlborough, MA, USA). Molecular weight and purity were determined by sodium dodecyl sulfate polyacrylamide gel electrophoresis (SDS-PAGE). Endotoxin contaminants were removed via Detoxi-Gel Endotoxin Removing Columns (ThermoFisher, Waltham, MA, USA). Final endotoxin content was determined to be below 0.1 EU/mL using Pierce™ LAL Chromogenic Endotoxin Quantitation Kit (ThermoFisher, Waltham, MA, USA).

#### Rat OA Model

Rats were anesthetized using 2.5% isoflurane (Patterson Veterinary, Greeley, CO, USA). Right knees were aseptically prepared with betadine surgical scrub (Purdue Products, Stamford, CT, USA) and 70% ethanol in triplicate, ending with a fourth betadine scrub. Via a 1-2 cm skin incision and blunt muscle dissection, the medial collateral ligament was exposed and then transected. Knee abduction was then used to expose the medial meniscus, with the meniscus transected radially in its central portion. Joint capsule and muscle were closed with absorbable 5-0 vicryl braided sutures (Ethicon, Somerville, NJ, USA). Skin was closed with 4-0 ethilon nylon monofilament (Ethicon, Somerville, NJ, USA). For post-surgical pain, rats received subcutaneous buprenorphine (0.05 mg/kg, Patterson Veterinary, Greeley, CO, USA) peri-operatively then every 12 hrs for 48 hrs.

#### Intra-articular Injections

Rats were anesthetized and aseptically prepared as described above. A 1 mL 27G x 3/8 syringe (Becton Dickinson, Franklin Lakes, NJ, USA) was inserted through the patellar ligament into the joint space. After injection, the needle was removed, gauze wetted with 70% ethanol was pressed against the injection site, and the knee was flexed.

#### IVIS Image Analysis

A region of interest (ROI) was drawn around the largest signal then copied to all other images. After subtracting background radiance, average radiance was calculated in the ROI. Average radiance for each animal was log_10_ normalized, plotted against time, then area under the curve (AUC) was calculated. Using AUCs, unpaired Student’s *t*-tests were conducted to evaluate clearance differences between NL and NL-Gal3 treated MCLT+MMT knees or NL and NL-Gal3 treated contralateral knees.

#### Gait Analysis

Rodent gait was analyzed using our GAITOR system (23). Briefly, high-speed videography (IDT M3, Pasadena, CA, 500 fps) and Kistler 3-axis load cells (±2 kN, Sindelfingen, Germany) were used to collect gait data during walking. Videos were analyzed to determine stride length, percentage stance time, temporal symmetry, and spatial symmetry (24). Force recordings were analyzed for peak vertical force at weeks 7-11 (25).

After IDO-Gal3 or saline treatment, gait differences were assessed with linear mixed effects models treating the animal identifier as a random factor. If indicated, comparison of least square group means was conducted using Tukey’s HSD corrections for multiple comparisons. For stride length, percentage stance time, and peak vertical force, data were first visualized relative to velocity, then statistically analyzed as both means and velocity-corrected residuals. For residualization, pre-treatment data (week 7) was used as the control line (24). For stance time imbalance and gait symmetries, data were also compared to mathematical definitions for balanced and symmetric gait using Bonferroni-correct Student’s *t*-tests.

#### Tactile Sensitivity

50% paw withdrawal thresholds were determined using Chaplan’s up-down method (26). Here, von Frey filaments (0.6, 1.4, 2.0, 4.0, 6.0, 8.0, 15.0, and 26.0 g, Stoelting Co., Wood Dale, IL, USA) were applied to the hind foot’s plantar region, starting with the 4.0 g filament. If paw withdrawal occurred, a less stiff filament was applied; if paw withdrawal did not occur, a stiffer filament was applied. Using these data, the force where paw withdrawal is equally likely to stimulus tolerance can be approximated (26). Tactile sensitivity was evaluated using a linear mixed effects model treating week and group as fixed effects and animal ID as a random effect. When indicated, comparisons of least squared means were conducted, correcting for compounding type 1 errors using Tukey’s HSD adjustment.

#### Magnetic Capture

Magnetic capture was used to assess intra-articular levels of IL6, CCL2, and CTXII (27). In magnetic capture, particles containing superparamagnetic iron oxide nanoparticles are used to magnetically precipitate target molecules from synovial fluid (28). In this work, biotin-streptavidin binding was used to functionalize magnetic particles (Dynabeads MyOne™ Streptavidin C1, Cat. #65001, Life Technologies, Carlsbad, CA, USA) with antibodies for CTXII (cat. #AC-08F1, ImmunoDiagnostic Systems, Copenhagen, Denmark), IL6, and CCL2 (cat. #517703 and 505908, Biolegend, San Diego, CA, USA). Antibody-functionalized particles were injected into rat knees, incubated for 2 hrs, then washed from the joint using five 50 μL PBS washes. Particles in wash solutions were magnetically separated then washed again. CTXII, IL6, and CCL2 were released from particles via a 15 mins treatment of 100 mM Glycine-Tris buffer (pH 3.1, containing 2% BSA and 2 mM EDTA). Following release, particles were magnetically separated, and supernatant was adjusted to pH 8.3.

In supernatants, CTXII was evaluated using the Cartilaps CTXII ELISA (cat. #AC-08F1, ImmunoDiagnostic Systems Cartilaps kit, Copenhagen, Denmark) according to manufacturer’s instructions. CCL2 was quantified using a rat CCL2 ELISA (Cat. # KRC1012 Life Technologies, Carlsbad, CA, USA) according to manufacturer’s instructions and buffer modifications described in Yarmola et al (29). IL6 was quantified using an ELISA developed from an antibody pair (biotin anti-rat IL6 detection antibody, cat. #517703, and purified anti-rat IL6 coating antibody, cat. #517701, Biolegend, San Diego, CA, USA). For this ELISA, coating antibodies were diluted in buffer containing 100 mM NaHCO3 and 34 mM Na2CO3 (pH 9.5) and incubated on Nunc MaxiSorp™ microwells (Cat. #434797, ThermoFisher Scientific Inc., Waltham, MA, USA) for 30 mins on a plate shaker then overnight (4°C). Plates were washed 5 times (PBS with 0.5% Tween-20), blocked for 1 hr in PBS containing 2% BSA and 10% heat-treated bovine serum (blocking solution), then washed again. Samples and standards were pre-incubated with detection antibodies for 30 mins at room temperature then overnight at 4°C, then added to the plate, incubated for 3 hrs, and washed. Avidin-HRP (100 μl, cat. #405103, Biolegend, San Diego, CA, USA) was diluted 500 times in blocking solution, incubated in the wells for 30 mins, and then washed. Finally, tetramethylbenzidine substrate (100 μL) was added and incubated for 15 mins, with the reaction stopped using diluted sulfuric acid (100 μL). Absorbance was then read at 450 and 650 nm.

To quantify particles, particles were re-suspended in 60 μL of PBS, sonicated (Model M1800H, Branson Ultrasonic Corporation, Danbury, CT, USA), and read for absorbance at 450 nm. Known particle concentrations were used as standards.

Following ELISA and particle quantification, biomarker levels were estimated as described in (28) with assay validation provided in (27,28). IL6, CCL2, and CTXII levels in rat knees were compared with one-way ANOVAs with *post hoc* Tukey’s HSD tests when indicated.

#### Histology

Knees were fixed in 10% neutral buffered formalin (Fisher Scientific, Pittsburgh, PA, USA) for 48 hrs, decalcified in Cal-Ex (Fisher Scientific, Pittsburgh, PA, USA) for 5 days, dehydrated using an ethanol ladder, and infiltrated with paraffin wax. Then, 10 μm frontal sections were acquired with at least one section taken at every 100 μm between the anterior to posterior meniscal horns. Slides were stained with toluidine blue; 3 sections per knee were assessed by 3 blinded graders using our histological grading software (GEKO) (30,31). Histological differences between IDO-Gal3 treated and saline treated knees were assessed using Student’s *t*-tests.

## Results

Galectin-3 (Gal3) increased local retention of NanoLuc™ Luciferase (NL, p<0.0001, Fig. 1A). At 4 weeks post-injection, NL-Gal3 was present on the femur, patella/fat pad, meniscus, and tibia of both knees, while NL alone was not present (Fig. 1B).

**Fig. 1:**
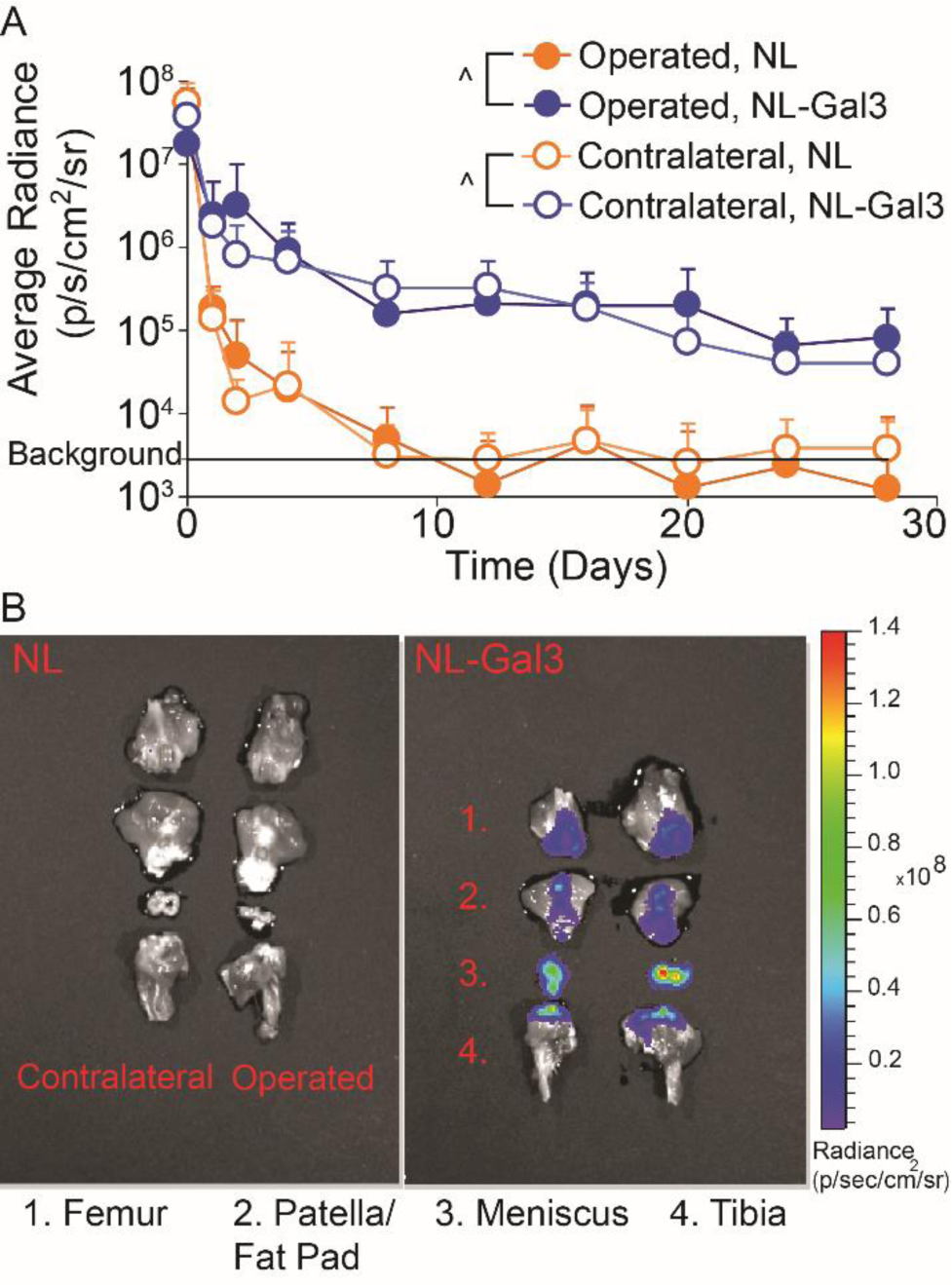
Joint retention and 4-week post-injection joint distribution for knees injected with NL and NL-Gal3. A) The joint residence of NL-Gal3 was significantly longer than the joint residence of unconjugated NL in both MCLT+MMT operated and contralateral joints (p<0.0001, linear mixed effects model). Data are presented as mean + 95% confidence interval. B) Post-mortem analysis of luminescence at 4 weeks after injection demonstrated the presence of NL-Gal3 on the femur, patella/fat pad, meniscus, and tibia of MCLT+MMT operated and contralateral joints. Unconjugated NL was not found.

The paw of MCLT+MMT-operated limbs became sensitized to touch after surgery, indicated by a decreasing 50% paw withdrawal threshold (p=0.002, Fig. 2A). After treatment with either saline or IDO-Gal3, tactile sensitivity improved relative to pre-treatment levels (p=0.0008), reaching the maximum of our testing range in 6 of 7 IDO-Gal3-treated animals. While the average tactile sensitivity for IDO-Gal3 and saline treated animals did not vary, increases in tactile sensitivity thresholds relative to baseline values were larger in IDO-Gal3 treated animals relative to saline treated animals at weeks 9-11 (p≤0.041, Fig. 2B).

**Fig. 2:**
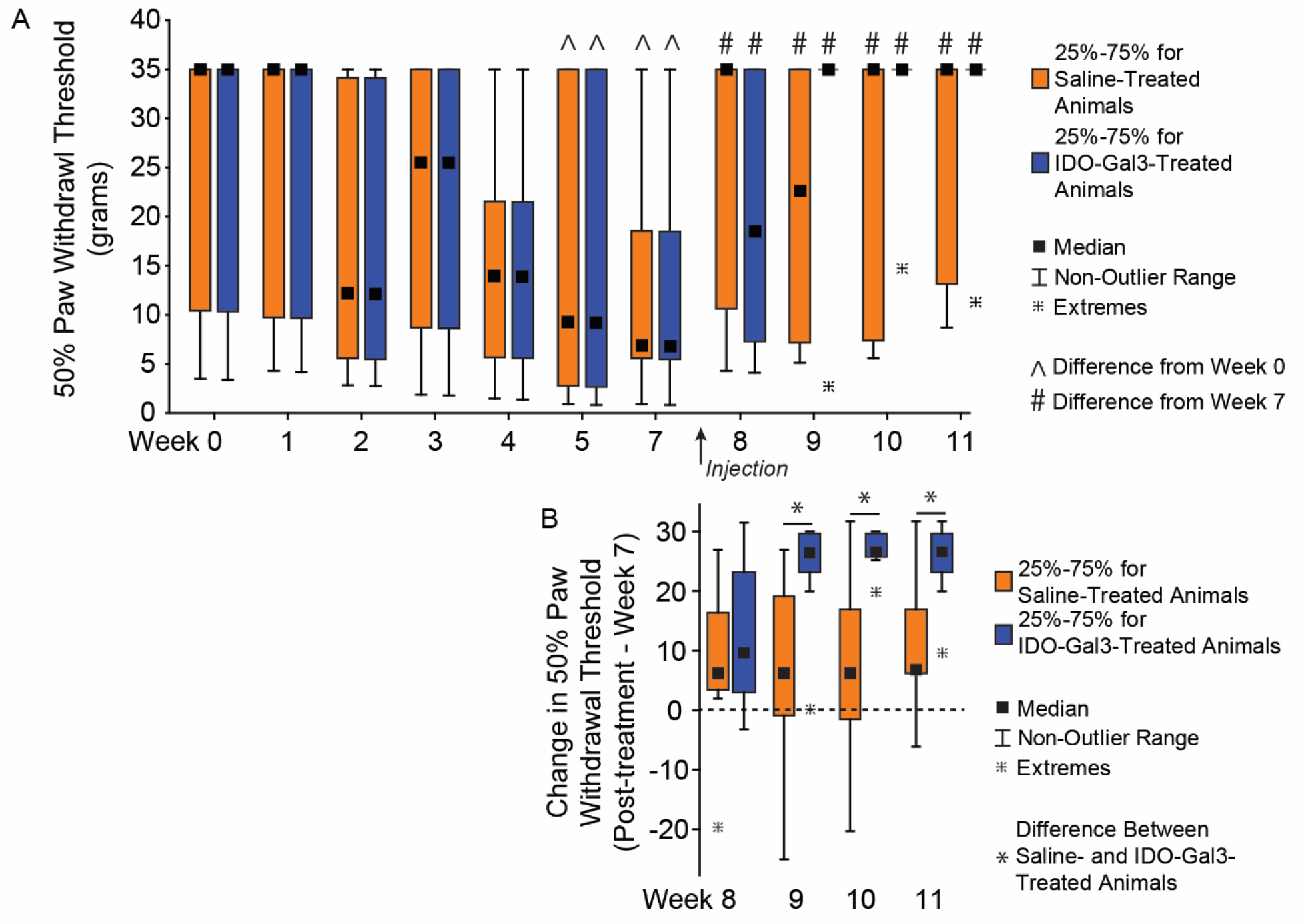
Tactile sensitivity before and after intra-articular injection of IDO-Gal3 or saline in MCLT+MMT operated knees. The 50% paw withdrawal threshold in MCLT+MMT operated limbs decreased at week 5 and week 7 relative to week 0 (^, p<0.05), indicating a heightened sensitivity to touch caused by MCLT+MMT surgery. Following intra-articular injections, 50% paw withdrawal thresholds increased in both saline and IDO-Gal3 treated animals relative to pretreatment (#, p<0.05), with 6 of 7 rats returning to baseline sensitivity levels in the IDO-Gal3 group by week 9. The average tactile sensitivity for IDO-Gal3 and saline treated animals did not vary after treatment; however, when change in tactile sensitivity relative to each animal’s week 7 value was assessed (paired analysis), increases in tactile sensitivity in IDO-Gal3 treated animals were larger than saline treated animals at weeks 9-11 (p≤0.041, Fig. 2B). As Chaplan’s up-down protocol provides a discrete set of data, results are shown as box plots with median, 25-75% interquartile range, and the lower and upper fence. Data outside the fence are plotted as individual data points.

Prior to treatment, saline and IDO-Gal3 treated rats used similar walking velocities (p≥0.89, Fig. 3A). Following treatment, IDO-Gal3 treated rats achieved higher velocities than saline controls on week 10 and week 11 (p<0.001, p=0.033, respectively). Before treatment, saline and IDO-Gal3 treated rats also used balanced, symmetric gaits. After treatment, the gait of saline treated rats became temporally asymmetric (asynchronous foot-strike sequence) at weeks 8-11, differing from IDO treatment animals at weeks 8-10 (p≤0.035, Fig. 3B). While neither saline nor IDO-Gal3 treated animals had imbalanced stance times, saline treated animals tended to spend more time on the contralateral limb (imbalance > 0), while IDO-Gal3 treated animals tended to spend more time on the affected limb (imbalance < 0). At weeks 8 and 9, stance time imbalance differed between saline and IDO-Gal3 treated animals (p≤0.025, Fig. 3C). Spatially asymmetric gait patterns were not observed in either group (Fig. 3D).

**Fig. 3:**
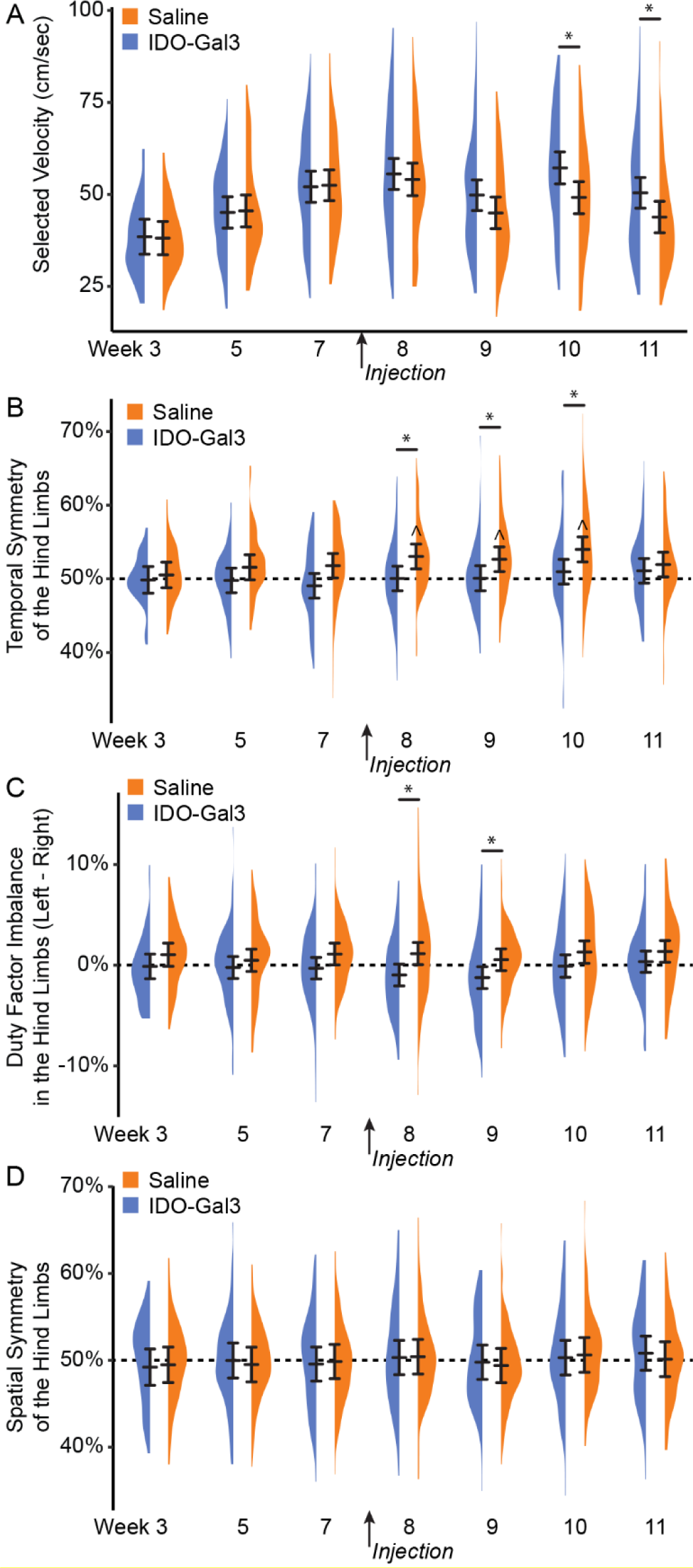
Walking velocity and spatiotemporal gait pattern analysis before and after intra-articular injection of IDO-Gal3 or saline in MCLT+MMT operated knees. Prior to treatment (weeks 3-7), saline and IDO-Gal3 treated rats used similar walking velocities (p≥0.89, Fig. 3A). However, following treatment, rats treated with IDO-Gal3 used faster walking velocities than saline controls on week 10 and week 11 (p<0.001, p=0.033, respectively). Balanced gaits are defined as stance time being equal on the left and right limb, or a difference of stance time between limbs being near zero. Temporally symmetric gaits have foot strike sequences that are equally spaced in time, where a right foot strike occurs halfway between two left foot strikes (i.e. temporal symmetry ≈ 0.5). Prior to treatment, saline and IDO-Gal3 treated rats used balanced, symmetric gaits. After treatment, the gait of saline treated rats became temporally asymmetric at weeks 8-11 (^symmetry > 0.5, p<0.05, Bonferroni-corrected *t*-test) and differed from IDO-Gal3 treatment at weeks 8-10 (p≤0.035, Tukey’s HSD pairwise test, Fig. 3B). Neither saline nor IDO-Gal3 treated animals had imbalanced stance times, though saline treated animals tended to spend more time on the contralateral limb (imbalance > 0) while IDO-Gal3 treated animals tended to spend more time on the affected limb (imbalance < 0). Here, at weeks 8 and 9, stance time imbalance differed between saline and IDO-Gal3 treated animals (p≤0.025, Tukey’s HSD pairwise test, Fig. 3C). Spatial symmetry measures the symmetry of the foot placement, rather than the timing of the foot strike. Again, a spatially symmetric gait has a right foot placement about halfway between two left foot placements (spatial symmetry ≈ 0.5). Here, both IDO-Gal3 and saline treated animals had spatially symmetric gait patterns throughout the experiment (Fig. 3D). This experiment includes over 1500 gait trials, with 65-131 trials collected at each treatment-timepoint. As such, data are plotted using density plots, with bars indicating the 95% confidence interval of each treatment-timepoint mean as predicted by our linear mixed effects statistical model.

Many gait parameters correlate to velocity (left column, Fig. 4). As such, velocity-dependent gait parameters were analyzed both as averages and velocity-corrected residuals, allowing for the evaluation of shifts due to and independent of velocity. In conjunction with velocity changes, differences between IDO-Gal3 and saline treated rats were found for average left (contralateral) limb percent stance time (p=0.04, treatment main effect), right (operated) limb percent stance time (p=0.005, week-treatment interaction), stride length (p=0.02, week-treatment interaction), and right (operated) limb peak vertical force (p=0.03, treatment main effect). Specifically, differences between IDO-Gal3 and saline treated rats were found for average left limb percent stance time at weeks 9, 10, and 11 (p<0.05), average stride length at weeks 9, 10, and 11 (p<0.05), and average right limb peak vertical force at weeks 9, 10, and 11 (p<0.04). Despite a significant interaction, differences at specific weeks were not detected for average right limb percent stance time. Peak vertical forces were similar in the left and right hind limbs of IDO-Gal3 treated rats, indicating balanced weight bearing. Saline treated rats reduced loads on the operated limb relative to their contralateral limb at weeks 9 and 11 (Supplemental Fig. 3). Again, to correct for velocity effects, residuals were calculated using week 7 data as the control line. Following velocity corrections, shifts in ground reaction forces were only significant in saline treated animals at week 9.

**Fig. 4:**
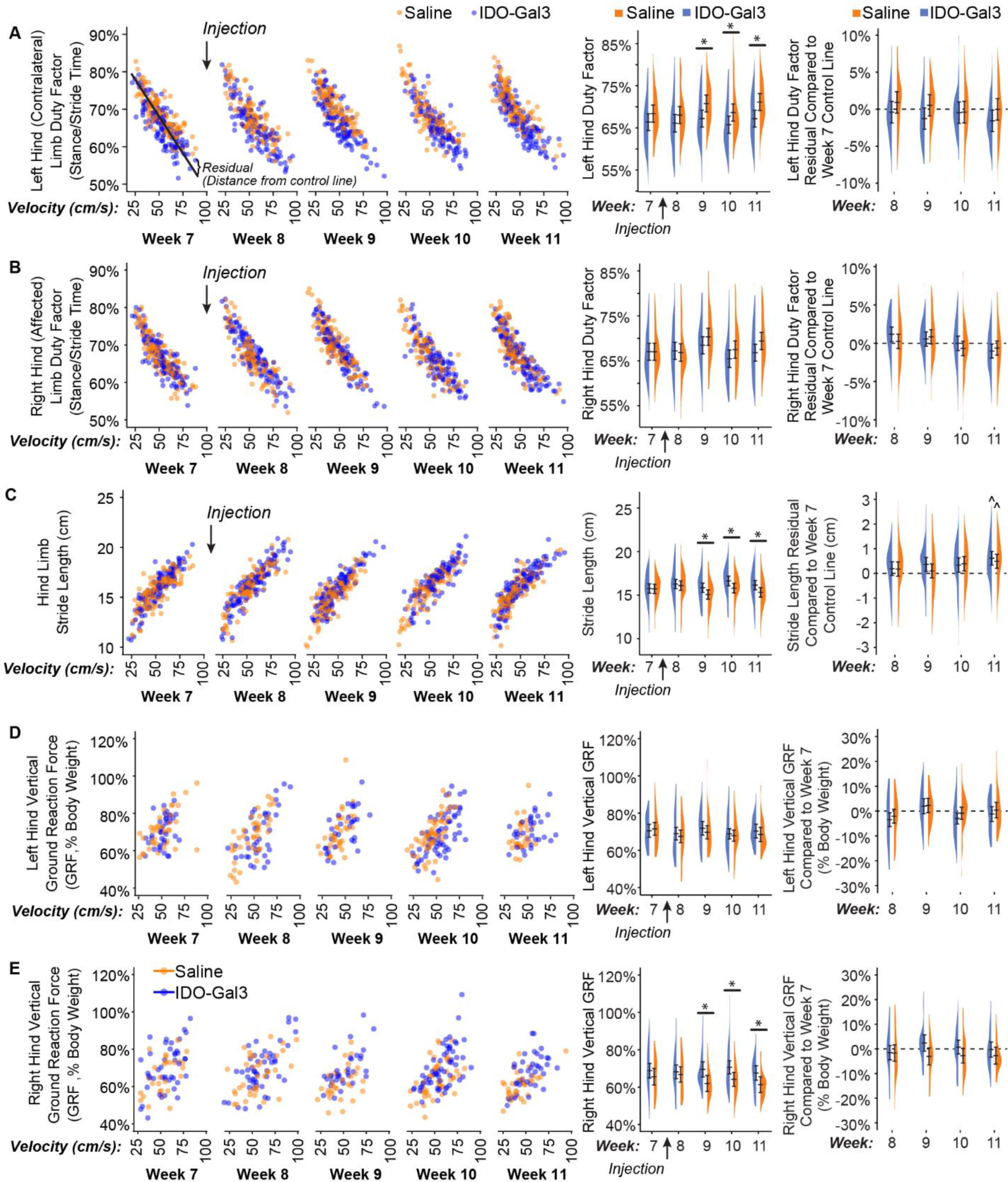
Gait measurements before and after intra-articular injection of IDO-Gal3 or saline in MCLT+MMT operated knees. Many gait parameters correlate to velocity. As such, velocity-dependent gait parameters are first shown relative to velocity (left column), with each data point shown. Then, shifts in gait parameters were analyzed both as averages and velocity-corrected residuals (using week 7 data as the control line). This allowed for the evaluation of gait shifts related to velocity changes (middle column) and independent of velocity changes (right column). In conjunction with velocity changes, differences between IDO-Gal3 and saline treated rats were found for average left contralateral limb percent stance time (p=0.04, treatment main effect), right operated limb percent stance time (p=0.005, week-treatment interaction), stride length (p=0.02, week-treatment interaction), and right operated limb peak vertical force (p=0.03, treatment main effect). Specifically, differences between IDO-Gal3 and saline treated rats were found for average left limb percent stance time at weeks 9, 10, and 11 (p≤0.048), average stride length at weeks 9, 10, and 11 (p≤0.046), and average right limb peak vertical force at weeks 9, 10, and 11 (p≤0.033). Despite a significant interaction, differences at specific weeks were not detected for average right limb percent stance time. Residualizing data to week 7 control lines eliminated these statistical differences, indicating differences in the gait patterns of IDO-Gal3 and saline treated rats were primarily related to increased walking speed. At week 11, stride lengths were longer in both IDO-Gal3 and saline treated rats relative to the week 7 control line (^, p<0.05); this is typical for experiments lasting longer than a month, as rats continue to have significant skeletal growth into adulthood. Again, data are plotted using density plots, with bars indicating the 95% confidence interval of each treatment-timepoint mean as predicted by our linear mixed effects statistical model.

At 4 weeks after intra-articular injections, interleukin-6 (IL6) levels in saline treated knees were elevated relative to contralateral controls (p=0.0025); moreover, IDO-Gal3 treated knees had lower IL6 levels relative to saline treated knees (p=0.0021, Fig. 5A). A similar trend occurred for intra-articular levels of CCL2 and CTXII, but neither CCL2 nor CTXII were statistically elevated in saline treated knees compared to contralateral controls (p=0.073, p=0.090, respectively). As such, shifts due to IDO-Gal3 treatment were also not statistically significant (Fig. 5B and 5C).

**Fig. 5:**
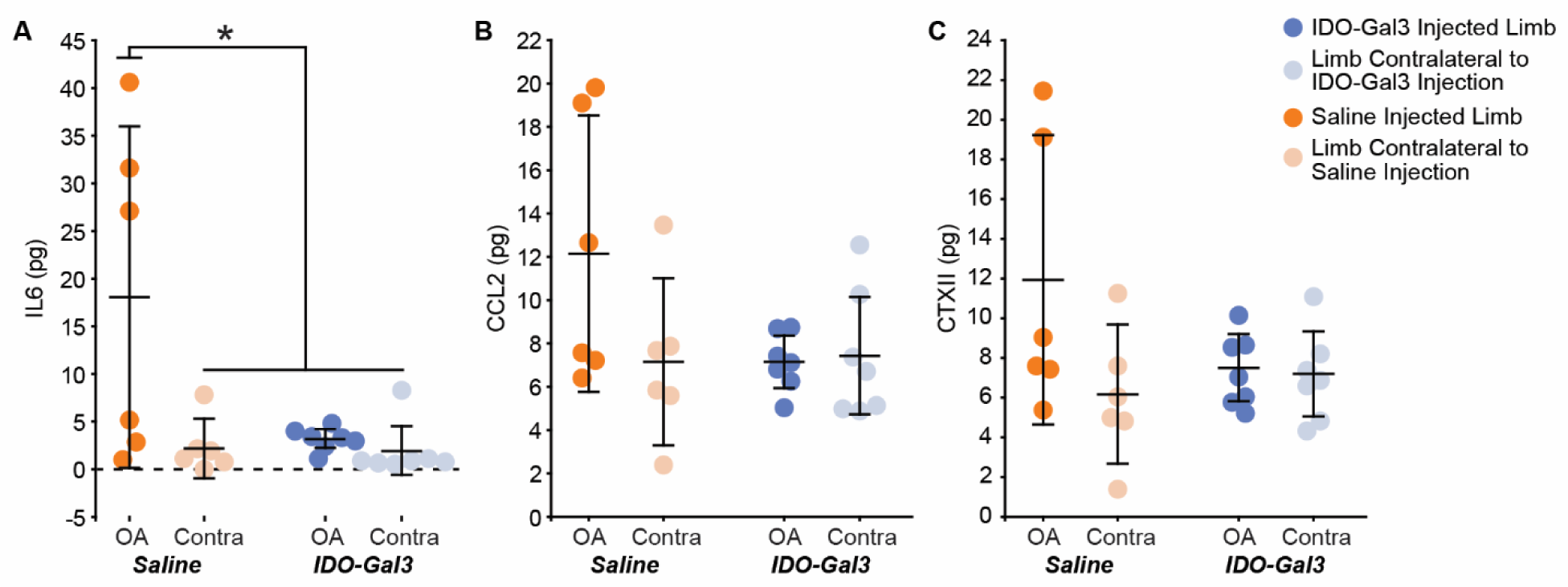
Intra-articular levels of IL6, CCL2, and CTXII 4 weeks after intra-articular injection of IDO-Gal3 or saline in MCLT+MMT operated knees. At 4 weeks after intra-articular injection, joint levels of IL6 were significantly decreased in the IDO-Gal3 treated operated knees compared to saline treated operated knees (A, p=0.03). Intra-articular levels of CCL2 (B) and CTXII (C) were not elevated in saline treated operated knees compared to contralateral controls, and as such, shifts due to IDO-Gal3 treatment could not be detected (p=0.073 for CCL2 and p=0.090 for CTXII). Bars represent mean +/− 95% confidence interval.

Please note, at the time of injection (8 weeks after MCLT+MMT surgery), full thickness cartilage lesions are expected (24). As such, all MCLT+MMT operated knees had marked cartilage damage. Treatment with IDO-Gal3 did not markedly alter cartilage histological measures, osteophyte size, or medial capsule thickness. However, IDO-Gal3 treated knees had lower epiphyseal trabecular bone area relative to saline controls (p=0.031, Fig. 6).

**Fig. 6:**
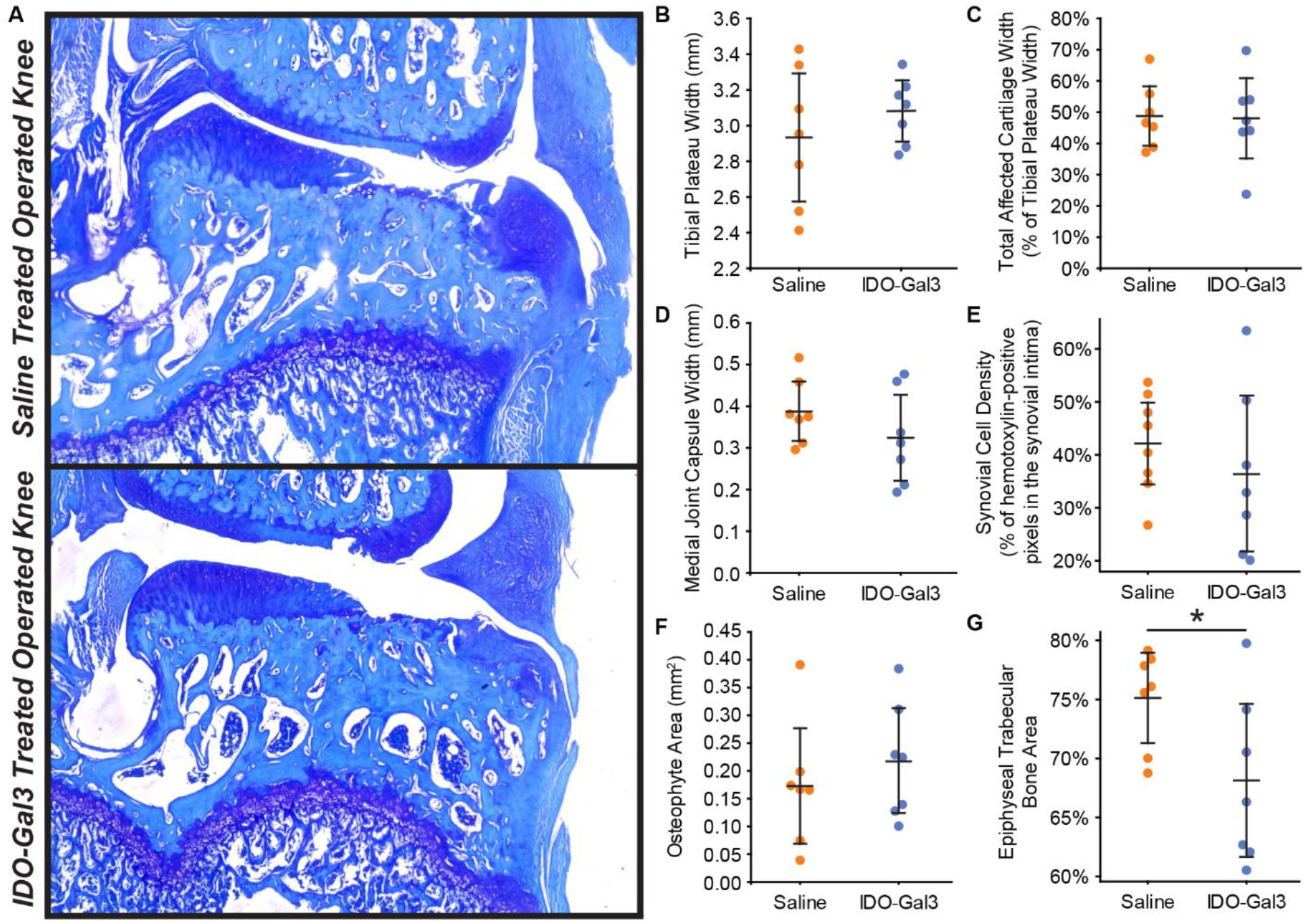
Histological scores of joint remodeling. At 12 weeks after surgery and 4 weeks after intra-articular injection, similar degrees of cartilage damage are seen in IDO-Gal3 and saline treated knees (A). In scoring the histological images, tibial plateau width (B), total affected cartilage width (C), medial joint capsule width (D), synovial cell density (E), and osteophyte area (F) were similar for IDO-Gal3 and saline treated groups. Epiphyseal trabecular bone area was lower in IDO-treated animals relative to saline controls, indicating marrow voids in the trabecular region were larger in IDO-Gal3 treated animals (G, p=0.031). Bars represent mean +/− 95% confidence intervals.

## Discussion

Inflammation is recognized as a key mediator of both joint destruction and OA pain (32,33). Thus, developing strategies for long-term control of inflammation remains a promising path for OA drug development. Many preclinical OA studies deliver therapeutics prior to or shortly after joint injury (34–37), when joint remodeling and pain-related behaviors are minimal. However, pain is the primary reason OA patients seek treatment (38). To this end, we developed a novel enzyme-based strategy for sustained control of joint inflammation and evaluated this therapeutic after behaviors indicative of OA pain had developed.

Our first goal was to demonstrate sustained intra-articular residence time of an enzyme due to Gal3 conjugation. Here, joint residence of nanoluciferase (a reporter enzyme) was extended by Gal3 conjugation relative to no conjugation. Gal3 has an affinity for β-galactosides found on extracellular matrix proteins, cell surface receptors, and lubricin (39–41). Gal3 also binds sulfated glycosaminoglycans, like chondroitin sulfate, which is abundantly found in healthy cartilage (21). However, in our data, operated and contralateral joints had similar nanoluciferase retention, suggesting Gal3 is not primarily binding to chondroitin sulfate in cartilage (which is lost as OA develops). Nonetheless, Gal3 conjugation extended joint residence in healthy and OA-affected joints, showing luminescence signal across multiple tissues at 4 weeks after injection in both healthy and OA-affected joints.

Use of Gal3 to extend joint residence requires careful consideration. Unconjugated Gal3 can have pro-inflammatory actions; however, these actions require that Gal3 can cluster cell surface receptors via an oligomerization that initiates cell adhesion, differentiation, or apoptotic signals (42). As shown in our prior work, conjugation of Gal3 to another protein prevents this action and limits these pro-inflammatory effects (22). Moreover, screens of synoviocyte viability in preparation for this study confirmed that IDO-Gal3 was not cytotoxic for the doses used in this study (Supplemental Fig. 4).

Our second goal was to evaluate IDO’s ability to modulate OA-associated inflammation. Here, IDO-Gal3 was injected into knees with established OA. At 4 weeks post-injection, IDO-Gal3 significantly decreased intra-articular IL6 levels. IDO-Gal3 tended to have similar effects on CCL2 and CTXII; however, in saline treated animals, neither CCL2 nor CTXII were significantly elevated in MCLT+MMT operated knees relative to contralateral controls. Thus, any effects related to IDO-Gal3 delivery were not detectable in our statistical analyses. While we have previously detected elevated CCL2 and CTXII levels in rodent OA models (27,29), this was in untreated animals and saline injection may have reduced effect sizes to some degree. Nonetheless, IDO-Gal3 did affect IL6 levels at 4 weeks after injection, demonstrating potential to provide sustained effects on OA-associated inflammation.

Our third goal was to evaluate IDO’s ability to modulate OA-related symptoms. Importantly, compensatory gaits can develop due to both pain avoidance and mechanical joint dysfunction, and in rodent OA models, these compensations can be unilateral (limping) or bilateral (shuffle-stepping), even for unilateral injury (24). Because both unilateral and bilateral compensations reduce loading on the injured limb (43), peak vertical force often provides the most sensitive gait variable for the evaluation of OA effects in rodents. Here, IDO-Gal3 treated rats had increased loading on the operated limb relative to saline treated animals at weeks 9, 10, and 11. Shifts in dynamic limb weight bearing were confirmed by spatial and temporal shifts in the gait cycle. For example, IDO-Gal3 treated rats also used symmetric and balanced gaits; saline treated animals were asymmetric and imbalanced at weeks 8 and 9 and asymmetric at week 10. IDO-Gal3 treated rats used higher walking velocities than saline treated animals at weeks 10 and 11. Finally, IDO-Gal3 treated animals used longer stride lengths than saline treated animals at weeks 9, 10, and 11. Combined, these data demonstrate improvement in limb use (as detected via gait parameters) in IDO-Gal3 treated animals relative to saline controls.

Gait parameters that are correlated to walking velocity were first shown as raw values, then analyzed as both average values and differences that were residualized to pre-treatment data. After accounting for velocity differences, the gait profiles of IDO-Gal3 and saline treated animals were no longer statistically significant. This indicates IDO-Gal3 treated animals did not use fundamentally different gaits, but instead, IDO-Gal3 treated animals achieved higher functional limb use (higher limb forces, longer stride lengths, improved stance times) via the use of faster walking velocities. Combined with the use of symmetric and balanced gaits, these data demonstrate a strong improvement in joint use for rats injected with IDO-Gal3.

Testing of tactile sensitivity evaluates withdrawal reflexes related to limb hypersensitivity. Here, IDO-Gal3 treatment substantially increased paw withdrawal thresholds relative to pre-treatment levels, with all but one IDO-Gal3 treated rat markedly increasing their paw withdrawal threshold after injection. Withdrawal thresholds also tended to increase with saline treatment, but to a lesser extent.

Histologically, MCLT+MMT surgery causes significant cartilage damage by 8 weeks post-surgery. Due to cartilage’s limited repair capacity, treatment at 8 weeks post-surgery was not anticipated to improve cartilage parameters. However, IDO-Gal3 treatment did alter IL6 levels, reduce tactile sensitivity, and improve gait parameters. Moreover, subchondral bone grades followed these changes in inflammation and behavior, where epiphyseal trabecular bone area indicated less area dedicated to trabecular bone and more area to marrow voids, trending toward percentages seen in healthy knees (approximately 60%, (31)). As such, it is plausible that treatment with IDO-Gal3 shortly after joint trauma could arrest OA progression; however, further studies are needed to address this question.

Several metabolic pathways are altered in OA joints, including tryptophan metabolism (44). IDO modulates the immune system primarily through local depletion of L-tryptophan and production of kynurenines (45). Recent work suggests IDO-driven shifts in local metabolites can cause immunomodulatory effects on T cells and antigen presenting cells (46). Consistent with these findings, the transfection of IDO plasmids into PMA-differentiated THP-1 (dTHP-1) cells results in increased expression of anti-inflammatory markers and decreased expression of pro-inflammatory markers (17). Due to inflammation, IDO is often upregulated in antigen presenting cells (45), and these antigen presenting cells can then activate regulatory T cells, inhibit effector T cell responses, and attenuate pro-inflammatory responses (47). Taken together, these qualities support IDO’s ability to restrain pro-inflammatory cascades. However, specific immunomodulatory effects of IDO in OA remain unanswered. Future work will also need to assess shifts in joint-level metabolic pathways and immune cell phenotypes after intra-articular IDO-Gal3 injection.

IDO has been explored in other forms of arthritis, primarily rheumatoid arthritis. Here, studies have focused on assessment of downstream metabolites, rather than behavioral assays. For example, IDO inhibition in collagen-induced arthritis (CIA) increased disease severity and elevated Th1 and Th17 responses in the joint (14,48). Related to this, intra-articular injection of dendritic cells modified to overexpress IDO reduced joint damage in the CIA model (49). However, inability to sustain IDO’s effects has limited progress (50). Our approach, directly injecting IDO conjugated to Gal3, may overcome these issues.

The limitations of the study include the loss of magnetic nanoparticles in a subset of our animals, lowering statistical power of our intra-articular inflammation analysis. Additionally, because intra-articular inflammation was prioritized in this study, the direct assessment of metabolites in synovial fluid was not possible. Furthermore, therapeutic efficacy was not evaluated as a function of dose or time after administration. We intend to evaluate these parameters in follow-up studies. Nonetheless, these data demonstrate proof-of-principle that intra-articular delivery of IDO-Gal3 affects the inflammatory environment in the OA joint for sustained periods of time and these effects correspond to changes in OA-related behaviors in animals with established knee OA.

## Conclusions

In this study, a novel enzyme-based therapeutic was tested in a rat model of established knee OA. First, we demonstrated that Gal3 conjugation can extend the joint residence of a reporter enzyme out to 4 weeks. Moreover, intra-articular injection of IDO-Gal3 in the OA-affected limb produced sustained effects on rodent walking, improved tactile sensitivity, and decreased IL6 levels. These data demonstrate the potential of IDO-Gal3 to serve as a metabolic reprogramming strategy for long-term, control of intra-articular inflammation and OA-related pain. Future studies will evaluate how intra-articular delivery of IDO-Gal3 modifies joint metabolism, immune cell polarization, and progression of joint damage in additional rat OA models.

## Author Contributions

BDP conducted and analyzed *in vitro* viability experiments, clearance and biodistribution experiments, tactile sensitivity experiments, gait analysis, and histological experiments. EBS expressed and purified IDO-Gal3. SAF expressed and purified NL-Gal3. EGY performed magnetic capture experiments and analyzed data with BDP. KDA performed MMT surgeries. BDP, EBS, BGK, GAH, and KDA contributed to the study design and data interpretation. The manuscript was prepared by BDP, with critical feedback from all authors. All authors agree with the content and interpretation of the data as presented.

## Acknowledgements

The authors would like to thank Natalie Thurlow and Mallory Box for assisting with gait processing. The authors would also like to acknowledge Nicole Sieling for sectioning and staining samples for histology, and Kiara Chan, Jacob Griffin, and Jessica Aldrich for grading histological images. We would also like to thank Sabrina Freeman for providing scientific and technical support regarding IDO.

---

**Supplemental Fig. 1:**
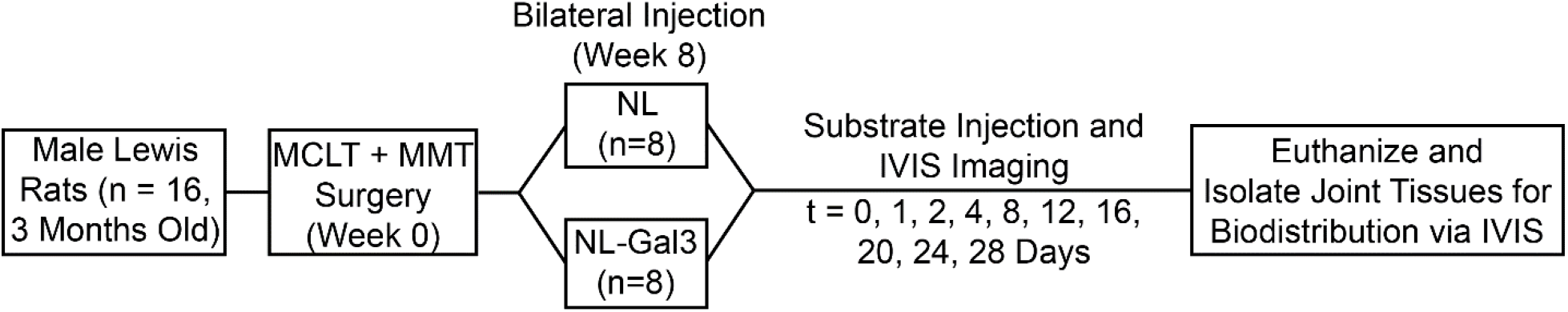
Flowchart describing the experimental design used to assess the joint residence and joint distribution of an enzyme after conjugation to galectin-3. Sixteen 3-month old, male Lewis Rats underwent medial collateral ligament transection plus medial meniscus transection (MCLT+MMT) surgery. At 8 weeks after MCLT+MMT surgery, rats were injected with either NanoLuc™ (NL, n=8) or NanoLuc™ galectin-3 (NL-Gal3, n=8) in both the operated and contralateral knees. Following NL or NL-Gal3 injection, rats were injected with the NL substrate, furimazine, and immediately imaged. Furimazine injections and IVIS imaging were repeated 1, 2, 4, 8, 12, 16, 20, 24, and 28 days after injection. After imaging on day 28, rats were euthanized and knees were dissected to isolate the patellar tissue, tibial tissue, femoral tissue, and meniscus for assessment of joint distribution.

**Supplemental Fig. 2:**
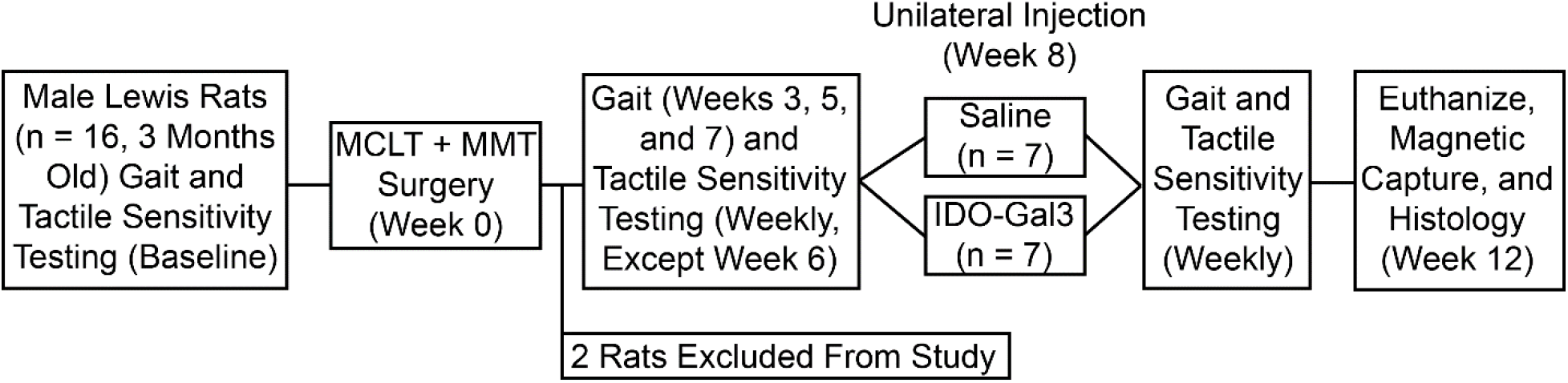
Flowchart describing the experimental design used to assess the ability of IDO-Gal3 to modulate OA-related pain and inflammation. Sixteen 3-month old, male Lewis rats underwent baseline gait and tactile sensitivity testing, then MCLT+MMT surgery. Following surgery, one rat was euthanized due to surgical complications, and another was excluded after histological evaluation showed no evidence of meniscus transection (OA did not develop). Remaining rats underwent gait assessments on post-surgery weeks 3, 5, 7, 8, 9, 10, and 11, and tactile sensitivity testing on post-surgery weeks 1, 2, 3, 4, 5, 7, 8, 9, 10, and 11. At 8 weeks after MCLT+MMT surgery, rats received either an injection of saline (n=7) or IDO-Gal3 (n=7) in the MCLT+MMT operated knee. At 12 weeks post-surgery, all rats were euthanized, then assessed for IL6, CCL2, and CTXII via magnetic capture and for joint damage using histological evaluations.

**Supplemental Fig. 3:**
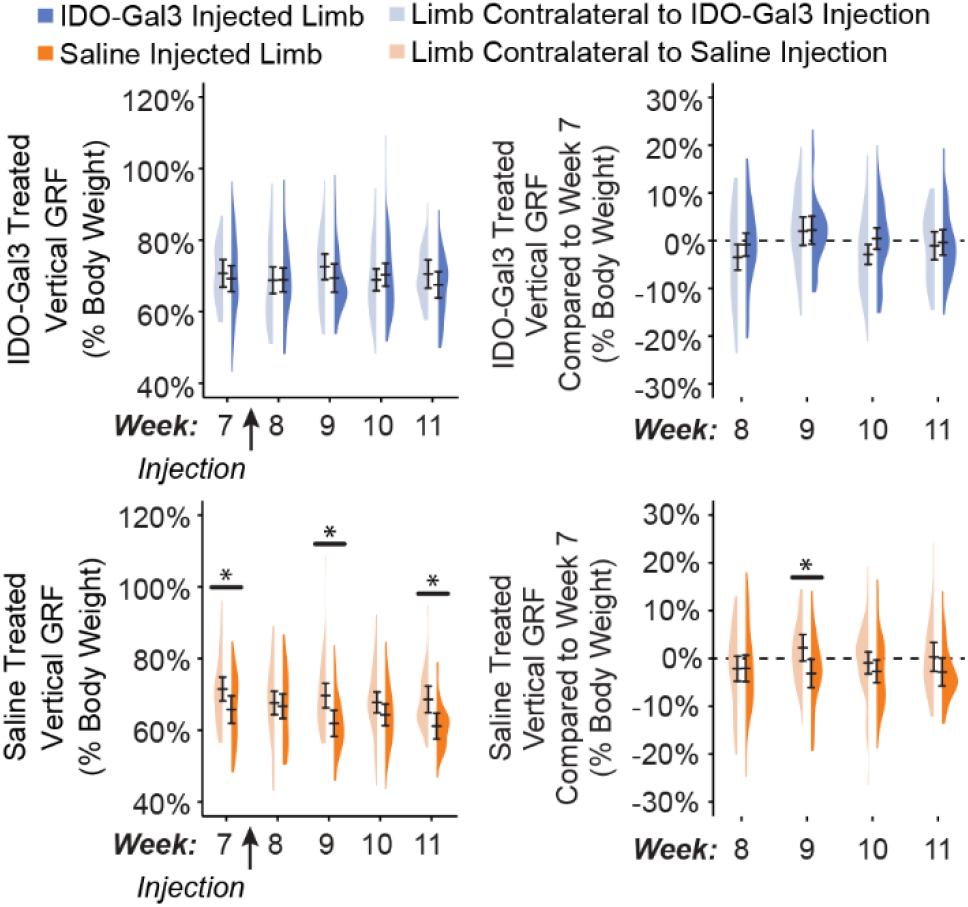
Peak vertical force measurements before and after intra-articular injection of IDO-Gal3 or saline into MCLT+MMT operated knees. This figure provides an alternate visualization of data presented in Fig. 4D and 4E, placing the left and right limbs of saline and IDO-Gal3 animals next to each other. Here, saline animals showed lower peak vertical forces in their affected limbs at week 9 and week 11 (p≤0.003). After residualizing data to the week 7 control line, differences at week 9 remained (p=0.009), while differences at week 11 were no longer significant.

**Supplemental Fig. 4:**
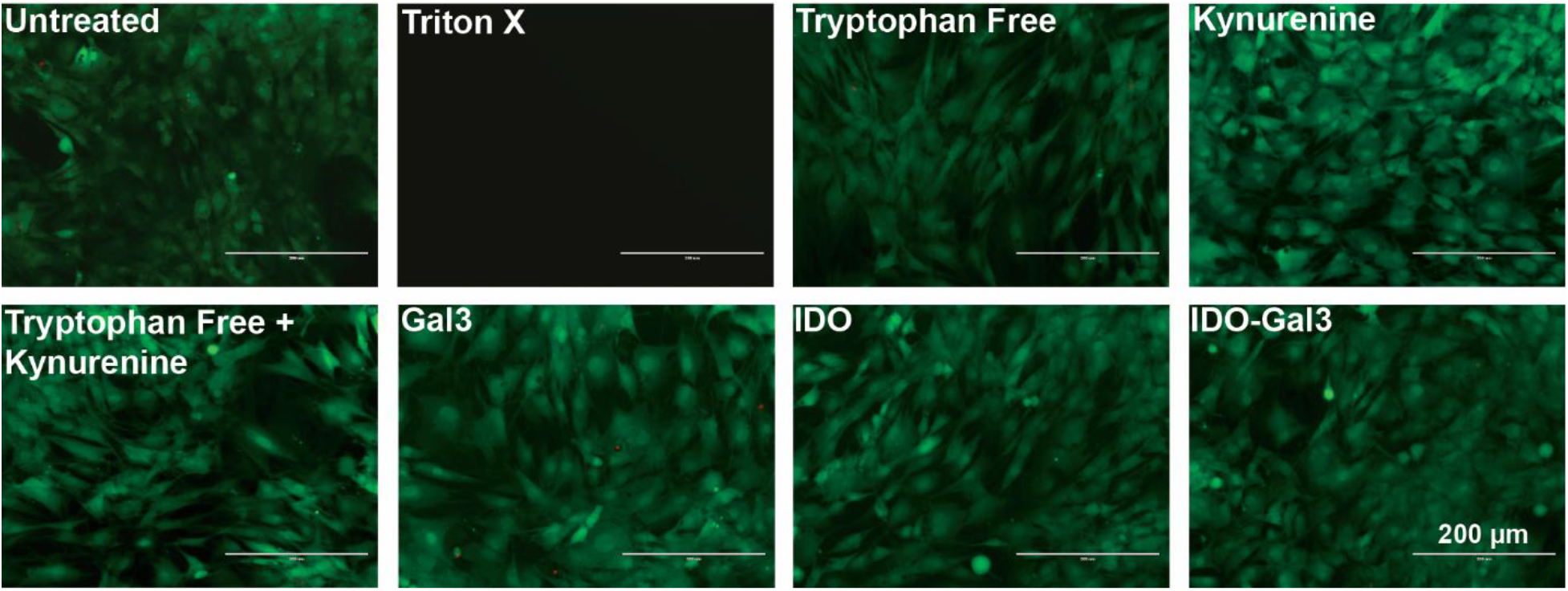
Synoviocyte viability after exposure to Gal3, IDO, and IDO-Gal3, as visualized with a LIVE/DEAD™ Cell Imaging Kit. In a preliminary experiment, rat synoviocyte viability after exposure to IDO-Gal3 was assessed via LIVE/DEAD™ Cell Imaging Kit (R37601, Invitrogen, Carlsbad, CA, USA) following manufacturer’s instructions. Cells were grown to 70%-80% confluence. Then, synoviocytes were incubated with Triton X-100, tryptophan free media, kynurenine-supplemented media (2080 μg/mL), tryptophan free media with a kynurenine supplement (2080 μg/mL), Gal3-supplemented media (8.3 μg/mL), IDO-supplemented media (15 μg/mL, and IDO-Gal3-supplemented media (2.37 μg/mL) (n=5/group) for 24 hrs. Concentrations of tryptophan and kynurenine were selected to be 100x greater than levels found in the OA joint (12.67 mM/mL and 8.74 μg/mL, respectively [1]); IDO concentrations were based on necessary levels to affect 100x concentrations of tryptophan in a 24 hrs period. Synoviocyte death was not observed in tryptophan free media, kynurenine-supplemented media, tryptophan free media with a kynurenine supplement, Gal3-supplemented media, IDO-supplemented media, and IDO-Gal3-supplemented media. Treatment with Triton X-100 (positive control) elicited complete cell death.

